# The marmoset default-mode network identified by deactivations in task-based fMRI studies

**DOI:** 10.1101/2023.08.28.555132

**Authors:** Audrey Dureux, Alessandro Zanini, David J. Schaeffer, Kevin Johnston, Kyle M. Gilbert, Stefan Everling

## Abstract

Understanding the default-mode network (DMN) in the common marmoset (Callithrix jacchus) has been challenging due to inconsistencies with human and marmoset DMNs. By analyzing task-negative activation in fMRI studies, we identified medial prefrontal cortical areas, rostral auditory areas, entorhinal cortex, posterior cingulate cortex area 31, hippocampus, hypothalamus, and basomedial amygdala as marmoset DMN components. Notable, medial and posterior parietal areas that were previously hypothesized to be part of the DMN were activated during visual task blocks. Seed analysis using resting-state fMRI showed strong connectivity between task-negative areas, and tracer data supported a structural network aligning with this functional DMN. These findings challenge previous definition of the marmoset DMN and reconcile many inconsistencies with the DMNs observed in humans, macaque monkeys, and even rodents. Overall, these results highlight the marmoset as a powerful model for DMN research, with potential implications for studying neuropsychiatric disorders where DMN activity and connectivity are altered.

## INTRODUCTION

The default-mode network (DMN) was initially identified as a large set of brain regions that were more active during baseline than task periods in several positron emission studies^1,2^. Numerous subsequent fMRI studies have confirmed the existence of a DMN in humans^3^, and its activation has been associated with various functions including mind wandering, social cognition, language and semantic memory, and the construction of a sense of self. Changes in DMN activity and connectivity have been found in several neuropsychiatric disorders, including Alzheimer’s disease, depression, schizophrenia, autism spectrum disorder, and attention deficit disorder ^3^.

Studies in rodents^3–5^ and nonhuman primates^6–9^ have also identified putative functional and anatomical homologues to the human DMN which promise to offer a window into investigating the DMN using targeted invasive recording and stimulation techniques. A small nonhuman primate that may hold tremendous potential for these studies is the common marmoset (*Callithrix jacchus*). This New World primate has a lissencephalic (smooth) cortex that is ideal for laminar neurophysiology and electrode array implantations^10^. Moreover, its small size allows for the usage of ultra-high field preclinical MR scanners^11^. However, the marmoset DMN seems to differ considerably from the human DMN, potentially limiting its translational value. While in humans the DMN consists primarily of the medial prefrontal cortex (mPFC), posterior cingulate cortex, anterior superior temporal cortex, middle temporal cortex, and angular gyrus, fMRI studies in marmosets have identified the dorsolateral PFC area 8, posterior parietal and posterior cingulate cortices as the DMN^12–18^. The identified marmoset DMN is puzzling for at least two reasons. First, the marmoset posterior parietal cortex exhibits saccade-related activity as identified by fMRI^19^ and single unit recordings^20^. Further, electrical microstimulation in posterior parietal areas LIP and MIP evokes contralateral saccades^21^. This is difficult to reconcile with it being part of the DMN. Second, the absence of mPFC areas in the marmoset DMN is surprising given that it has been identified as a main component of the human DMN^1,3^ and is also part of the DMN in Old World macaque monkeys^7,9^ and even in rodents^4,5,22^. For these reasons, we revisited the search for the marmoset DMN by analyzing task-negative activation in our recently published fMRI studies in marmosets^23–26^.

## RESULTS

Here we aimed to isolate the marmoset’s DMN by identifying brain areas that exhibited higher activations during baseline period than task epochs, i.e., areas that decreased their activation during task blocks, in our previous marmoset fMRI studies. All studies employed passive tasks, i.e., the marmosets were simply presented with a set of videos and/or audio clips. During baseline periods, monkeys were presented with a small, filled black circle on a grey screen. Please note that we restrict our analysis to task-positive versus task-negative activations here and ignore all differences between the different conditions, which we already described in previous publications.

Since the human DMN was initially identified by areas that were deactivated during attention-demanding tasks^1,2^. we first checked whether we could identify the typical DMN in human subjects during such a passive task. To this end, we re-examined task-positive versus task-negative activations in data from our recent study^23^ in which we presented theory-of-mind and random Frith-Happé animations to 10 humans and 6 marmoset monkeys. Figure 1, top shows the task-positive (orange and red colors) and the task-negative activations (blue color) for human subjects in this task. Consistent with numerous human studies, we found task-negative activations in medial prefrontal cortex (including areas 25, 10v, 10d, p32, d32, p32pr, 9m, p24, 24dv), posterior cingulate cortex (23ab), superior temporal sulcus (STSda, STSdp), and angular gyrus (PFm, PG). The task-positive network included mainly visual cortical areas (e.g. V1, V2, V3, V4, MT, MST, FST, PH), posterior parietal cortex (areas PF, LIPv, 7PC, 7M, PCV), and premotor and frontal eye fields (area 6a, FEF, 55b, PEF, IFJp, 6r). These findings demonstrate that these passive tasks are able to evoke robust DMN activations in human subjects through task-negative responses.

**Figure 1.**
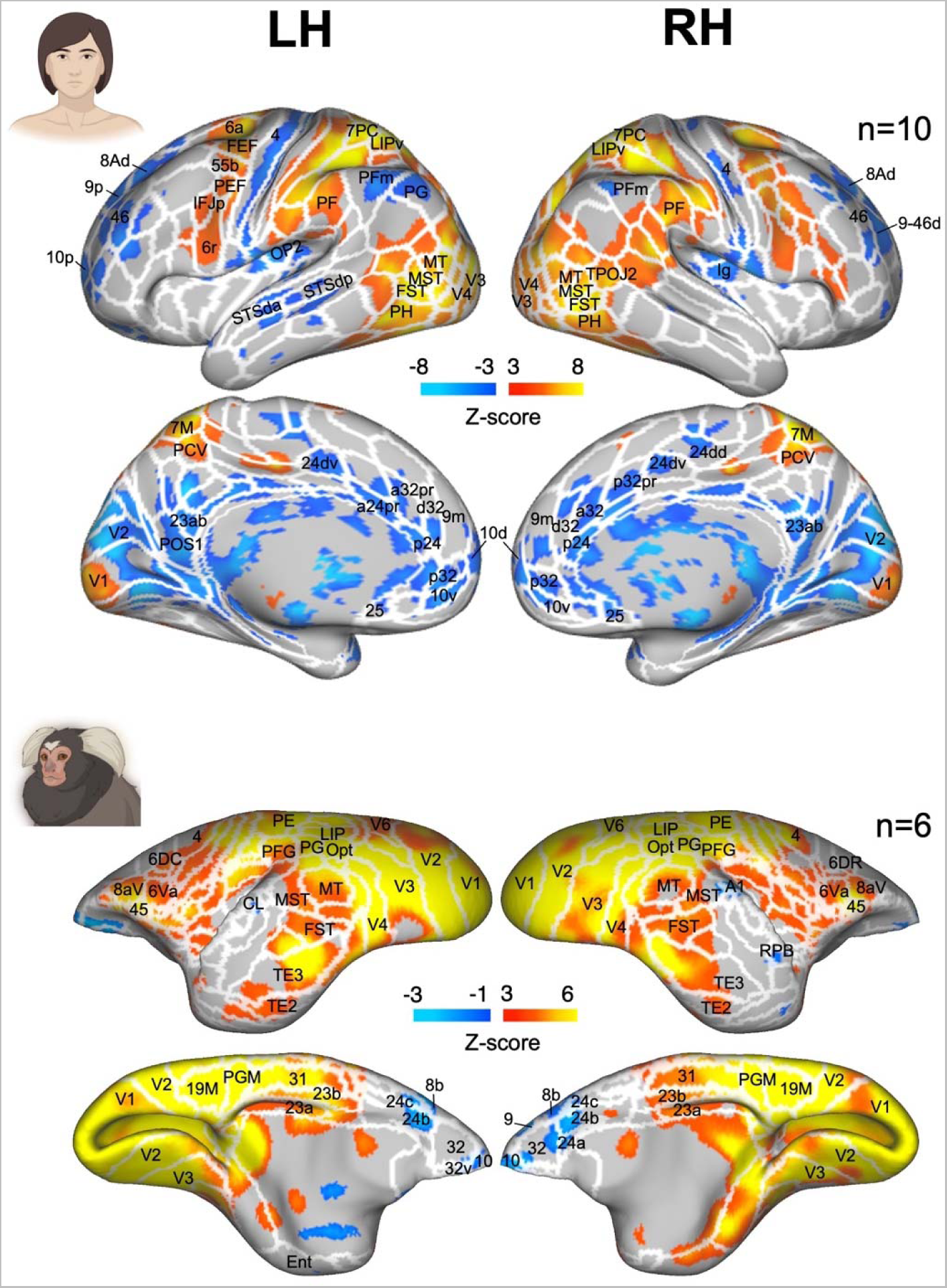
Task-positive and task-negative activations from our recent theory-of-mind and random Frith-Happé animations study (Dureux et al., 2023c). Group functional maps showing significant task-positive and task-negative activations, comparing all Theory-of-mind (ToM) and Random animations against baseline. Maps are displayed on right and left fiducial (lateral and medial views) for both human (top) and marmoset (bottom) cortical surfaces. The human map encompasses data from 10 participants, and the marmoset map includes 6 subjects. The demarcating white line highlights regionsas as per the recent multi-modal cortical parcellation atlas ^27^ for humans and based on the Paxinos parcellation of the NIH marmoset brain atlas ^28^. The brain areas reporting task-positive activation have threshold corresponding to z-scores >3 with yellow/orange scale, whereas the brain areas reporting task-negative activations have threshold of z-scores < −1 or −3, depending on the species, with blue scale (AFNI’s 3dttest++).

Next, we compared the task-positive and task-negative activations for the same task in marmosets (Figure 1, bottom). On the medial surface, task-positive areas included V1, V2, V3, 19M, PGM, 31, 23a, and 23b. On the lateral surface, activations were observed in V1, V2, V3, V4, V6, 19DI, Opt, MT, FST, MST, LIP, MIP, PG, and TE3. In addition to these widespread posterior activations, we also found activations in frontal areas 6DR, 8aV, and 6Va, and observed activations in somatomotor areas. Conversely, we found task-negative activations, i.e., deactivations during the task, in medial prefrontal areas 10, 32, 32v, 24a, 24b, 24c, 9, and 8b, as well as in the right primary auditory cortex and rostral parabelt (RPB). In contrast to humans, there were no task-negative activations in posterior cingulate cortex, or in area PG.

We then re-analyzed five additional of our recently published and in preparation fMRI studies by pooling all task epochs together and comparing them to the baseline periods (Fig. 2, left side).

**Figure 2.**
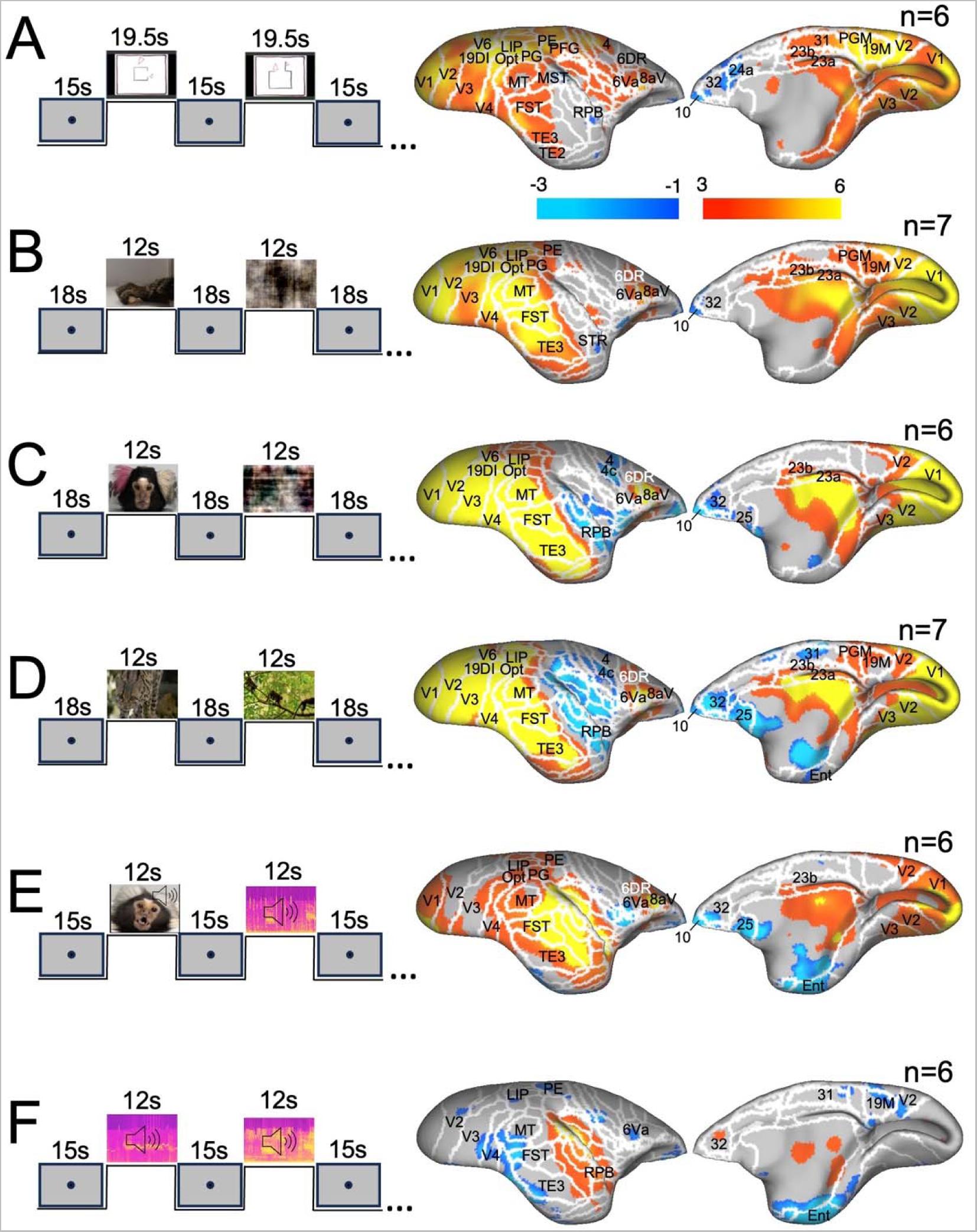
Task-related activations from 6 distinct tasks used in our recently published and in preparation fMRI studies. The left side of the figure outlines the task design for each experiment and the right side illustrates brain regions indicating positive or negative activations. This contrasts all task conditions against baseline periods for each study. In Studies A-D, marmosets were exposed to video stimuli without auditory components. Study E combined both visual and auditory stimuli, and Study F was exclusively auditory. Specifically, tasks included: TOM and Random Frith-Happé animations (A)(REF); videos of marmoset hands and forearms reaching towards objects or empty locations and their scrambled versions (B)(Zanini et al., 2023); neutral and negative marmoset face videos alongside their scrambled versions (C) (Dureux et al., 2023a); videos of predators or marmosets and their scrambled counterparts (D, unpublished data); unimodal auditory stimuli comprising vocal and scrambled calls, video stimuli with marmoset faces and scrambled versions, and combined audiovideo variants (E)(REF); and marmoset vocalizations and their scrambled equivalents (F, unpublished data). Group maps, varying from 6 to 7 marmosets depending on the task, are depicted on the right fiducial marmoset cortical surface, covering both lateral and medial views. The white line delineates the regions based on the Paxinos parcellation of the NIH marmoset brain atlas ^28^. Task-positive activation regions are tied to z-scores > 3 (yellow/orange scale), and task-negative zones correspond to z-scores < −1 (blue scale) as processed by AFNI’s 3dttest++.

In four of our studies (Fig. 2A-D), marmosets were presented with different types of videos without any audio. These included in addition to the Frith-Happé animations (Fig. 2A), videos of marmoset hands and forearms reaching towards objects or empty locations and their scrambled versions (Fig. 2B)^26^, videos of neutral and negative marmoset faces and their scrambled versions (Fig. 2C)^24^, and videos of predators or marmosets and their scrambled versions (Fig. 1D, unpublished data). Overall, these four studies showed very similar task-positive activations in visual, parietal, and lateral frontal regions. Although always considerably weaker than the task-positive activations, we also observed a clear pattern in task-negative activations. All tasks were associated with task-negative activations in medial prefrontal areas 10 and 32. Notably, two of the studies (Fig. 2 B and D) also showed task-negative activations in area 25. In addition, all four tasks were associated with some task-negative activations in temporal auditory areas. In the Frith-Happé animation study^23^, deactivations were restricted to the rostral parabelt (RPB) while in the action observation study^26^, they were localized to the superior temporal rostral (STR) area (Fig. 2A, B). In contrast, the other two studies^24^ showed strong task-negative activations across multiple auditory areas (Fig. 2C-D).

In addition, these two studies also exhibited task-negative activations in primary motor cortex, similar to the results in humans (Fig. 1, top).

We also included a study in which we presented auditory stimuli (vocal and scrambled calls), videos (marmoset faces and scrambled faces), and videos with corresponding audio (faces with calls, and scrambled faces with scrambled calls) (Fig. 2E)^25^. In this study, fewer visual areas were activated by the task conditions, but we observed very strong activations in auditory cortices. In addition, we also found frontal task-positive activations in area 6DR, 6Va, and 8aV. In contrast, task-negative activations were present in medial prefrontal areas 10, 32, and 25. A purely auditory study (marmoset calls and scrambled calls) elicited task-related activations in auditory cortices including core, belt, and parabelt areas (Fig. 2F). In addition, area 32 was activated. Task-negative activations were more scattered and included area 6Va, parts of V3 and V4.

Furthermore, three studies (Fig. 2D-F) also exhibited task-negative activations in entorhinal cortex (Ent) and posterior cingulate area 31.

To identify areas that were activated during multiple studies, we computed probability maps for task-positive and task-negative activations (Fig. 3). The analysis shows that the task-positive network included predominately areas in the occipital, parietal, and inferotemporal cortex (Fig. 3A). In addition, frontal area 6Va, 6DR, and 8aV were activated by most tasks. At the subcortical level (Fig. 3B), we observed consistent recruitment of the lateral pulvinar (Pul), lateral geniculate nucleus (LGN), lateral amygdala (LA), and the caudate nucleus (CN).

**Figure 3.**
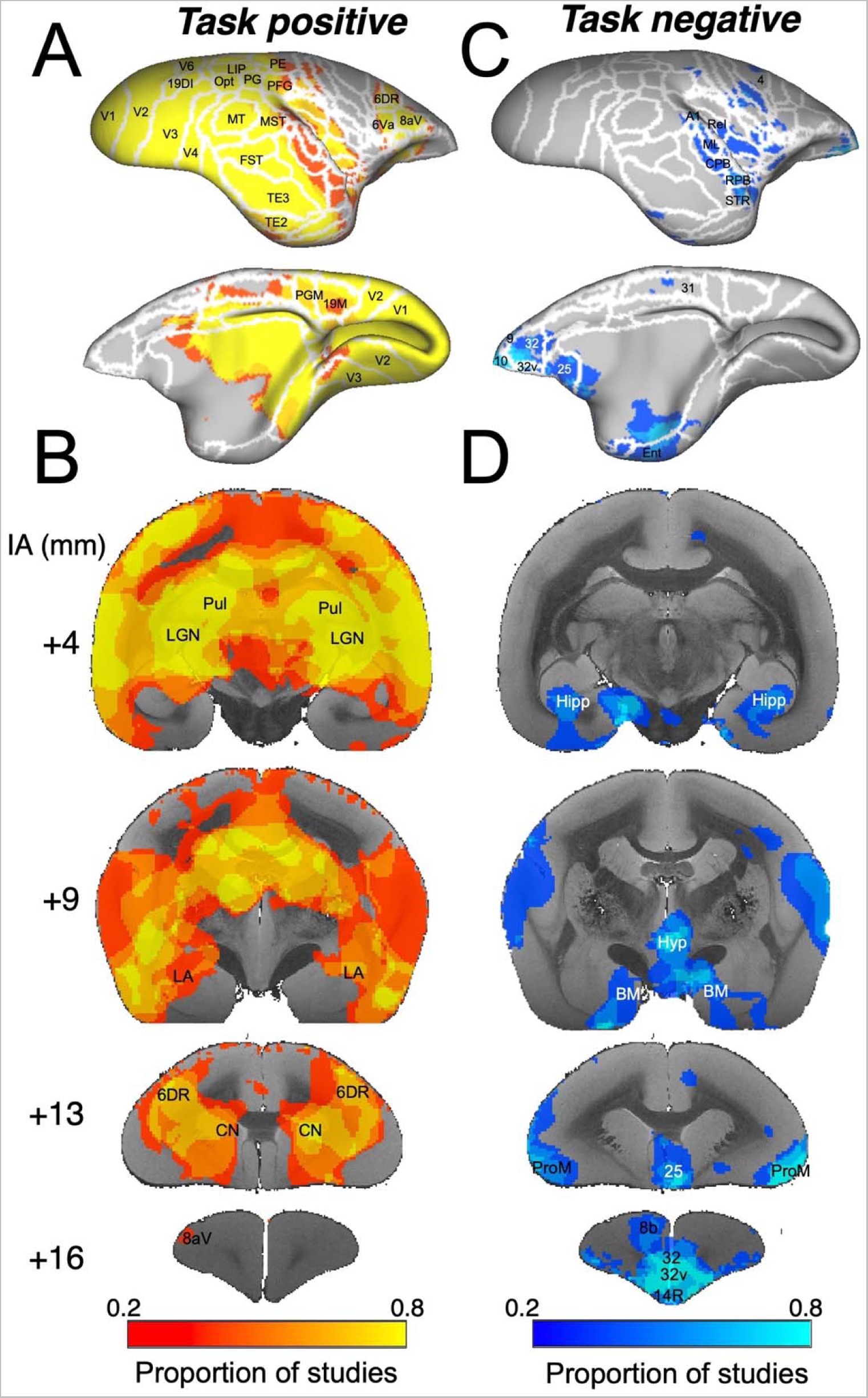
Probabilistic functional mappings of task-positive (A and B) and task-negative (C and D) activations. We generated coincidence maps of both task-positive and task-negative activations with a set threshold of z=3 for task-positive and z=-1 for task-negative activations. These were then transformed into probability maps using a custom Matlab program. The resulting maps are displayed on the left fiducial brain surfaces, showcasing both lateral and medial views (A and C), as well as on coronal sections (B and D). Distinct brain regions are highlighted based on the NIH marmoset brain atlas (Liu et al., 2018).

The task negative network included at the cortical level predominately medial prefrontal area 10, 32, 32v, and 25 (Fig. 3C). Further, primary auditory cortex (A1), caudal parabelt (CPB), RPB, and area STR displayed task-negative activations. In addition, entorhinal cortex (Ent) and area 31 were sometimes part of task-negative areas. At the subcortical level (Fig. 3D), the hippocampus (Hipp), basal medial amygdala (BM), and hypothalamus (Hyp) showed task-negative activations in several of our studies.

The hypothalamus has recently been identified as a core component of the human DMN^29^. To further investigate its functional role in the task-negative and task-positive networks, we performed a seed-based functional connectivity analysis of the hypothalamus using resting-state fMRI data from an open dataset resource^30^. Figure 4 shows the results of positive functional connectivity with the regions around the paraventricular nucleus of the hypothalamus in blue and negative functional connectivity in red. The functional connectivity maps showed positive correlations with auditory core, belt, and parabelt areas, predominately towards anterior locations, and with medial prefrontal areas 10, 32, 32v, and 25. At the subcortical level we found positive functional connectivity with the medial pulvinar (mPul) and hippocampus (Hipp). Overall, the positive functional connectivity of the hypothalamus resembled the task-negative network and the negative functional connectivity included most of the areas of the task-positive network.

**Figure 4.**
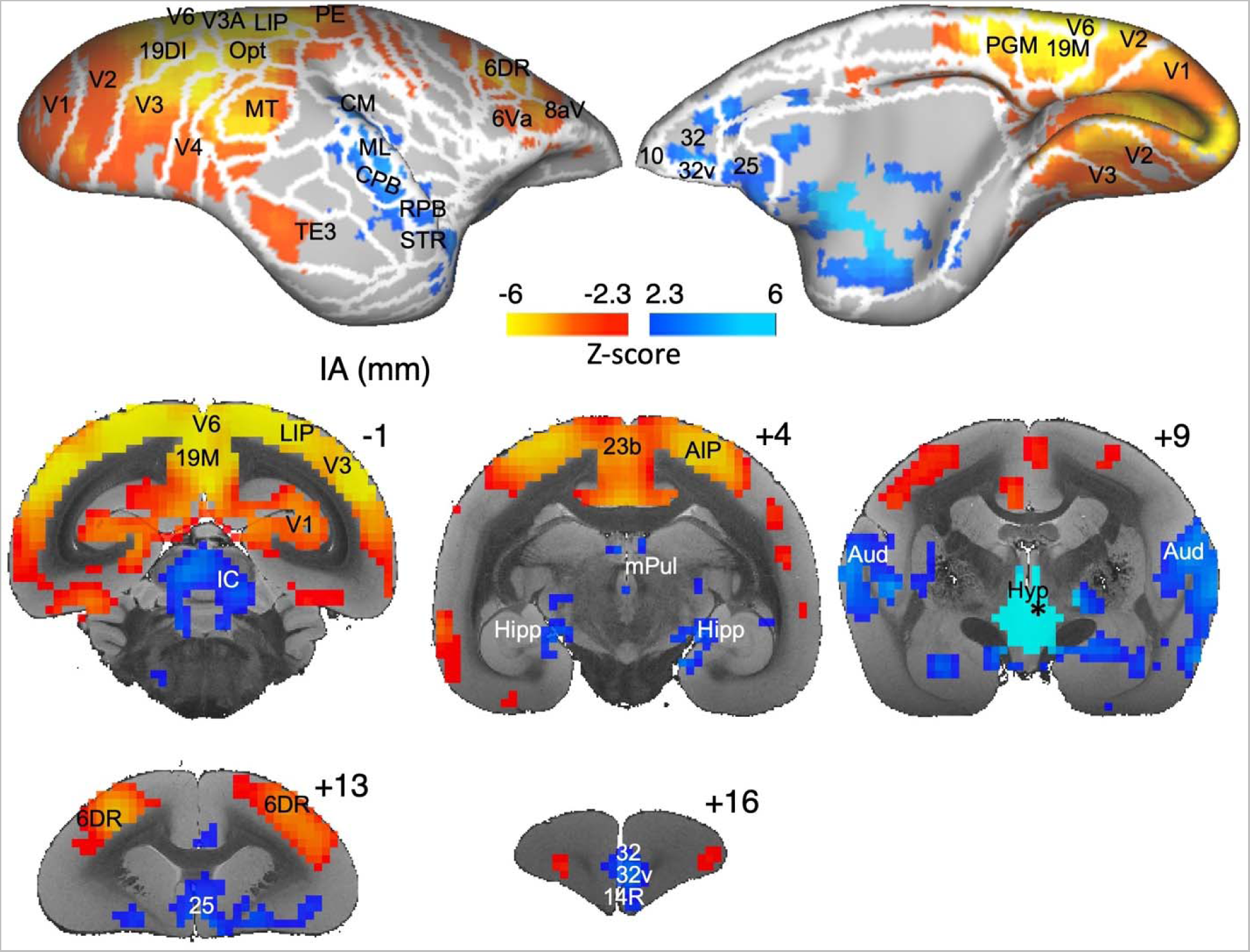
Functional connectivity of the hypothalamus. Seed-based functional connectivity map centered on the hypothalamus, particularly the regions around the paraventricular nucleus, derived from resting-state fMRI data (https://marmosetbrainconnectome.org) (Schaeffer et al., 2022a). Positive functional connectivity is represented in blue (z-scores > 2.3), while negative functional connectivity is shown in orange (z-scores < −2.3). This functional connectivity representation is superimposed on the left fiducial brain surfaces maps (both lateral and medial views) using the NIH marmoset brain template and atlas (Liu et al., 2018) and is also overlaid on coronal sections.

To test whether we would find a similar pattern of positive functional connectivity with other task-negative areas and negative functional connectivity with task-positive areas, we displayed functional connectivity maps of areas 32, 10, 25, RPB, hypothalamus, hippocampus, and basomedial amgydala on flatmaps of the right hemisphere (Fig. 5). These maps show strong functional connectivity (blue) with the areas of the task-negative network and strong anticorrelations (orange and yellow) with areas of the task-positive network.

**Figure 5.**
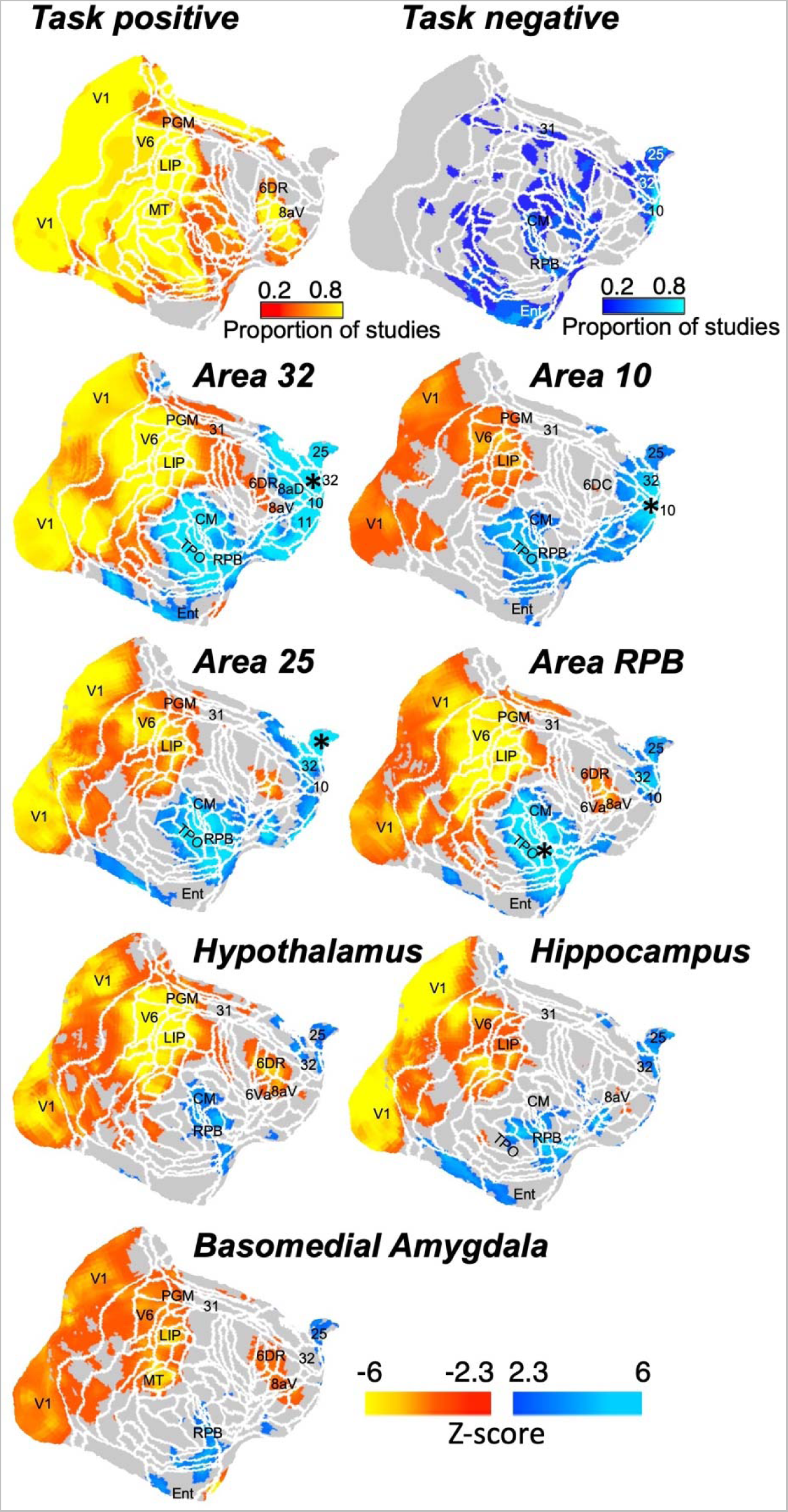
Functional connectivity maps of areas 32, 10, 25, RPB, hypothalamus, hippocampus, and basomedial amygdala. The figure illustrates a seed-based functional connectivity map focused on areas 32, 10, 25, RPB, hypothalamus, hippocampus, and basomedial amygdala, as derived from resting-state fMRI data (https://marmosetbrainconnectome.org) (Schaeffer et al., 2022a). Positive functional connectivity is depicted in blue (with z-scores > 2.3), while negative functional connectivity is rendered in orange (with z-scores < −2.3) and displayed on flat maps of the right hemisphere. At the top of the figure, to assess if a similar pattern of positive functional connectivity with other task-negative areas and negative functional connectivity with task-negative areas can be observed, we displayed the probabilistic functional mappings maps described in figure 3 on flat maps.

With evidence from functional connectivity^30^ (Fig. 5) that marmoset anterior cingulate (namely area 32/32v) is not connected with posterior cingulate (areas 23 and 31), we hypothesized that the structural underpinnings of these connections would also be sparse^31^. Moreover, examining the functional connectivity across the entire medial and lateral frontal cortices^32^ with posterior cingulate, we posit that the lateral frontal cortex (namely area 8) has much stronger connectivity to areas 23 and 31 than do any of the anterior cingulate regions. To validate this hypothesis, we conducted a comparison between the anatomical connections of the region bordering mPFC areas 32 and 32v, and those of lateral PFC area 8aD. This comparison utilized viral anterograde tracer data openly accessible through the BRAIN/MINDS resource^33^. Consistent with our proposed hypothesis, the findings from this examination (illustrated in Fig. 6) reveal a clear anatomical linkage between the lateral PFC (area 8aD) and the posterior cingulate regions 23 and 31. In contrast, the connectivity pattern of the mPFC area 32/32v is notably focused within the mPFC itself, extending to include the orbitofrontal areas 47O and 13, as well as temporal areas TPO, TE1, and STR. Additionally, there are weak projections from more posterior part of area 29 to the mPFC. In this pattern of results, the absence of structural connectivity between the mPFC and the posterior cingulate cortex stands out in the marmoset.

**Figure 6.**
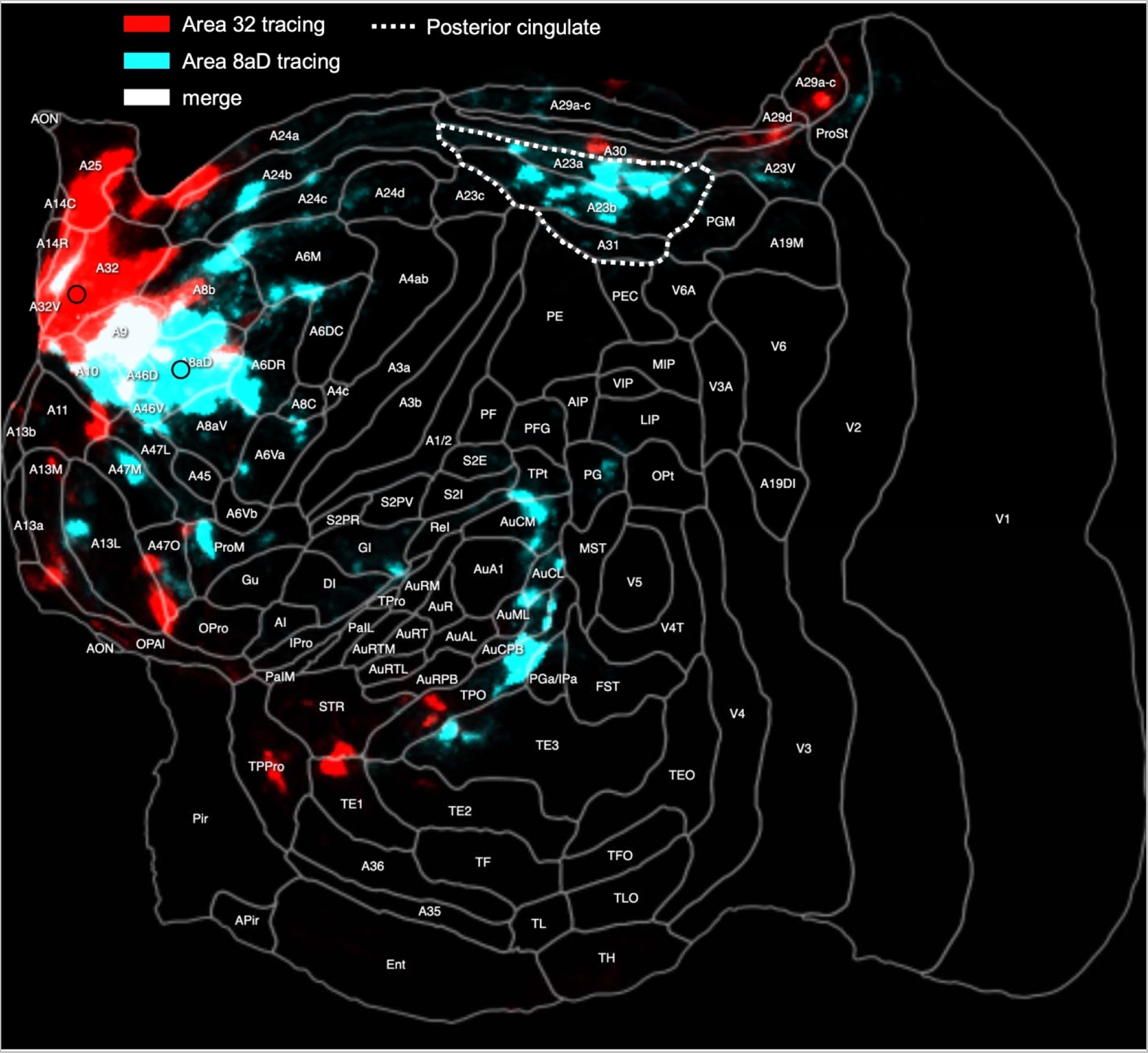
Structural connectivity of the medial prefrontal cortex area 32/32v and lateral prefrontal cortex area 8aD. Viral anterograde tracer data from the BRAIN/MINDS publicly available marmoset tracer resource^33^. In red, anatomical connectivity of the region bordering mPFC areas 32 and 32v; in light blue, anatomical connectivity of the lateral PFC area 8aD; in white their overlap. The dashed white line indicates the localization and extent of the posterior cingulate region, encompassing areas 23 and 31.

## DISCUSSION

Our findings revise the structure and potential function of the default-mode network (DMN) in the common marmoset (*Callithrix jacchus*). By focusing on task-negative activations from our previously published tasks, we discovered that the DMN in marmosets incorporates the medial prefrontal cortex (mPFC), parts of the auditory cortex, hippocampus, basomedial amygdala and the hypothalamus. This was particularly the case for studies where we presented visual stimuli during task blocks, whereas vocal stimuli activated some of these areas during the task blocks. Importantly, single neuron activity in area 32 of the marmoset mPFC that is activated by vocal stimuli (see also Fig. 1F), also decreased during baseline blocks, suggesting a robust contribution to the DMN.

Most studies to date have relied on resting-state fMRI to identify the DMN in marmosets^12,15–17^. Importantly, these studies have emphasized the presence of the posterior cingulate cortex/precuneus in the putative marmoset DMN using independent component analysis (ICA), or they have even identified the putative DMN by placing a seed in the marmoset posterior cingulate cortex. However, the DMN in humans was initially defined as a set of brain regions that showed higher activations during baseline than task epochs^1,2^ and not as areas that showed a particular functional connectivity pattern. Liu and colleagues^14^ used a similar approach to what we employed here, and attempted to identify the marmoset DMN as a network of brain regions that increased their activity during the baseline period in an fMRI task using visual stimuli. By using this approach, they found activations in V2, 19M, PGM, 23V, V6A, PEC, LIP, VIP, MIP, PG and OPT during the baseline period. They did not detect any activations in prefrontal cortex. This stands in sharp contrast to the absence of any activations in lateral or medial parietal areas during baseline blocks in our analysis of visual blocked fMRI tasks. In fact, lateral and medial parietal areas were activated by all videos. We believe that this difference relates to the specifics of the task data that Liu et al. (2019) analyzed in their study. In their task^14,34^, the monkeys were presented with 16 s of visual pictures of faces, body parts, objects, and scrambled stimuli, during which they had to maintain fixation within a 5-degree central circle to receive a liquid reward every 1.5s. In contrast, no fixation requirements were imposed, and no reward was given during the 20 s baseline blocks (grey screen). Therefore, one can assume that the marmosets generated more and larger saccades during the baseline periods than during the task blocks in this specific design (see also Fig. 2A of Hung et al. for example eye traces^34^). Consequently, the activations in the posterior parietal cortex observed by Liu and colleagues^14^ during the baseline period are likely related to an increase in saccadic eye movements during the baseline period, consistent with a role of the marmoset posterior parietal cortex in saccade control as shown by electrical microstimulation^21^, electrophysiological recordings^20^, and fMRI^19^. In contrast, the task-based fMRI studies that we used here to identify the DMN had no fixation requirements.

The discovery of a prominent role of mPFC in the marmoset DMN reconciles the previous discrepancies observed with humans^3,17^, macaque monkeys^6,7,9^, and even rodents^4,5,22^ in which mPFC areas have been identified as part of the DMN. We also found activations in three of the six studies in posterior cingulate area 31. However, consistent with Liu and colleagues^14^, we did not find any functional connectivity between the medial prefrontal areas and posterior cingulate cortex in marmosets using rs-fMRI data. Accordingly, tracer data from the Brain/MINDS Marmoset Connectivity Resource^33^ confirm monosynaptic connections between area 32 and other regions of the task-negative network such as areas prefrontal areas 25, 9, 10 and 14, the most frontal part of area 24a, the temporoparietal (TPO) area, and STR, only weak monosynaptic connections with the posterior cingulate region. The absence of this human hallmark DMN connectivity pattern points to a specific functional change in DMN structure between humans, old world macaques, and marmosets^17^.

The inclusion of parts of the auditory and anterior temporal cortex in the marmoset DMN opens novel research avenues concerning auditory processing and its integration with default mode functioning. In fact, the DMN that we identified showed strong overlap with the vocal network that we recently found by auditory fMRI in awake marmosets^35^. In humans, the anterior temporal cortex and the middle temporal cortex are integral parts of the DMN^3^. These regions contain vocal patches^36–39^ and are also parts of the human language network^40^. Based on the observation in a macaque PET study that showed higher activations predominately in mPFC, auditory cortex, and insular cortex during rest versus the performance of a visual working memory task^6^, Watanabe and colleagues hypothesized that the default mode activity may reflect some form of thought in the monkeys^6,8^. Our findings in marmosets resemble these findings in macaques and the observation that regions such as mPFC area 32 and parts of auditory cortex are more active during baseline than task blocks is intriguing, given that we have recently shown that these regions are also activated by the processing of conspecific vocalizations^35^. Therefore, it will be particularly interesting to investigate the functional interactions between mPFC areas and auditory areas in marmosets, even in the absence of external auditory stimuli or tasks.

At the subcortical level, we observed task-negative activations in the basomedial amygdala consistent with its role in the human DMN^29,41,42^. Interestingly, in mice, this region has been identified as a major target of the mPFC^43^. Within the basomedial amygdala of mice, neurons can differentiate between safe and threatening environments, with their activation— mediated by projections from the mPFC—helping to alleviate high-anxiety states. In marmosets, mPFC area 25 has been associated with cardiovascular and behavioural responses to stress^44^, and both anterograde and retrograde tracer data demonstrated a direct connection between marmoset mPFC and the basomedial amygdala^45^.

The identification of the hypothalamus as a prominent subcortical component of the marmoset DMN supports a recent human DMN model^29^. Interestingly, resting-state fMRI from a large open resource marmoset dataset^30^ confirmed positive functional connectivity between the hypothalamus and other task-negative areas and negative functional connectivity with the task positive network. Moreover, seed-based resting-state fMRI functional connectivity analysis showed that the different nodes of the task-negative network exhibited strong functional connectivity with each other and exhibited negative functional connectivity with regions of the task-positive network. This is completely consistent with previous findings in humans^46,47^. In accordance with this pattern of functional connectivity, data from studies using both anterograde and retrograde tracer methods have demonstrated a monosynaptic connection between the hypothalamus and the mPFC in marmosets^45^.

Overall, our results show that a network of structurally and functionally connected cortical and subcortical brain regions exhibits increased activity during baseline blocks particularly in visual blocked fMRI tasks. These findings demonstrate that the common marmoset is a promising nonhuman primate model for research into the neural processes within the DMN, with potential implications for understanding neuropsychiatric disorders where DMN activity and connectivity show marked changes.

## METHODS

For this study, we reanalyzed our previously published and in preparation studies using task-related awake marmoset fMRI^23–26^, and one task in humans^23^. Full experimental details are provided in the published studies.

### Marmosets

All experimental procedures were conducted in accordance with the Canadian Council of Animal Care policy and were approved by the Animal Care Committee of the University of Western Ontario Council on Animal Care. Additionally, the procedures complied with the Animal Research: Reporting In Vivo Experiments guidelines.

For the fMRI experiments, a total of 11 adult common marmosets (*Callithrix jacchus*) served as subjects. The number of animals utilized in each study ranged from 6 to 7, and both male and female marmosets were included.

All marmosets were pair-housed at 24 - 26° C with 40-70% humidity under a 12 h light-dark cycle.

### Preparation of the animals

Animals were implanted for head-fixed fMRI experiments with an MR-compatible head restraint chamber^48^ or with a machined PEEK (polyetheretherketone) head post^49^ under anesthesia and aseptic conditions as previously described^48,50^. Marmosets were acclimatized to the head-fixation system in a mock MRI environment over a three week training period^49^ that included head restraint and exposure to pre-recorded MRI sounds .

### Human participants

Ten healthy individuals (3 females, aged between 27 and 44 years) participated in the study^23^. Each participant was right-handed, had either normal vision or vision that was corrected to normal, and did not have any neurological or psychiatric disorders. The Ethics Committee of the University of Western Ontario approved the study, and all participants furnished written consent for their involvement.

### fMRI experimental setup

Visual stimuli were projected onto a forward-facing plastic screen positioned at a viewing distance of 119 cm using an LCSD-projector (Model VLP-FE40, Sony Corporation, Tokyo, Japan) via a back-reflection on a first surface mirror. We used Keynote software (version 12.0, Apple Incorporated, CA) for stimulus display. The onset of visual and/or auditory stimuli was synchronized with an MRI TTL pulse triggered by a python program running on a Raspberry Pi (model 3B+, Raspberry Pi Foundation, Cambridge, UK). During each experimental run, different conditions (movies and/or auditory stimuli) were presented during task blocks. The duration of the task blocks varied between 12-19.5s between the studies but was constant within each study. The task blocks were interleaved by baseline blocks that varied between studies from 15-18s, during which a central black dot was displayed in the center of the screen against a gray background. Marmosets were not required to fixate in any of the studies.

### MRI data acquisition

All imaging studies were performed at the Center for Functional and Metabolic Mapping at the University of Western Ontario. For marmosets, the data were collected using a 9.4T/31 cm horizontal bore magnet and a Bruker BioSpec Avance III console running the Paravision 7 software package. We used a custom-built 15-cm inner diameter gradient coil (Handler et al., 2020)with a maximum gradient strength of 1.5 mT/m/A. Depending on the experiment, it was coupled with either a five receive channels^24,26^ facilitated by a head restraint chamber, or eight receive channels^23–26^ utilizing a machined PEEK head post. To ensure optimal signal reception, preamplifiers were located behind the animal. The receives coil was placed inside an quadrature birdcage coil of 12-cm inner diameter, which was custom-fabricated in-house and served for transmission. Gradient-echo based single-shot echo-planar images (EPI) were acquired with specifications of 0.5 mm^3^ isotropic resolution, 42 slices [axial], 400 kHz bandwidth, and a GRAPPA acceleration factor of 2 (left-right). The repetition time (TR) was 1.5s for all purely visual studies, In auditory experiments, every slice also acquired within 1.5 s, succeeded by a 1.5s period in which the scanner noise was low. During one of the sessions, a T2-weighted structural image was collected for each animal with the following parameters: TR=7s, TE=52ms, field of view=51.2x51.2 mm, resolution of 0.133x0.133x0.5 mm, number of slices= 45 [axial], bandwidth=50 kHz, GRAPPA acceleration factor: 2. The total experimental time was usually around 60 minutes per animal and included experimental setup, animal preparation, and scanning time.

For the human subjects, the imaging was conducted on a 7T/68 cm horizontal bore magnet (Siemens Magnetom 7T MRI Plus). This was combined with an AC-84 Mark II gradient coil, an in-house developed 8-channel parallel transmit, and a 32-channel receive coil^51^. Multi-Band EPI BOLD sequences were acquired with the following parameters: TR = 1.5s, TE = 20ms, flip angle = 30°, field of view=208x208 mm, matrix size = 104x104, resolution of 2 mm3 isotropic, number of slices= 62, GRAPPA acceleration factor: 3 (anterior-posterior), multi-band acceleration factor: 2. Additionally, field map images were also computed, derived from the core magnitude image and its corresponding phase images. For every participant, an MP2RAGE structural image was acquired during the sessions with the following parameters: TR=6s, TE=2.13 ms, TI1 / TI2 = 800 / 2700 ms, field of view=240x240 mm, matrix size= 320x320, resolution of 0.75 mm3 isotropic, number of slices= 45, GRAPPA acceleration factor (anterior posterior): 3.

### MRI preprocessing

The preprocessing of data from marmoset subjects was performed using AFNI^52^ and FSL^53^ software. From the anatomical images of each marmoset, a T2-weighted template mask was generated. This data was reoriented and subsequently, a manually skull-stripped mask was created via FSLeyes. The mask was then binarized using AFNI’s 3dcalc function. The anatomical data was then merged with the binarized mask to generate the T2 mask. Finally this mask was aligned with the 3D NIH marmoset brain atlas (NIH-MBA)^28^.

The raw functional data was converted to NIfTI format using dcm2niix. After this conversion, data was reoriented and corrected for any motion artifacts through functions like fslswapdim, fslroi, and topup. The dataset for each specific run was then interpolated using the applytopup function. Outliers that emerged were identified with 3dToutcount and subsequently eliminated using 3dDespike.

Thereafter, time-shifting was initiated using 3dTshift. For each run, the median volume served as the foundational base for alignment through 3dvolreg. Spatial smoothing was performed by applying a three-dimensional Gaussian function with a full-width-half-maximum (FWHM) ranging between 1 and 2 mm, depending on the study. To conclude the preprocessing phase, the frequency domain was restricted between 0.01-0.1 Hz using a band-pass filter, specifically the 3dbandpass function.

Data preprocessing for human subjects was conducted using SPM12 (Welcome Department of Cognitive Neurology, ^54^). The process began by converting raw images into NifTI format. Once transformed, functional images were subjected to field map correction using the specific toolbox in SPM, which drew upon both magnitude and phase images. Following this, the functional images were realigned to account for head movements and subsequently underwent slice timing correction. Both the anatomical and functionally corrected volumes were then coregistered with the MP2RAGE structural scan specific to each participant, facilitating their normalization to the Montreal Neurological Institute (MNI) standard brain space. Anatomical images were meticulously segmented to distinguish white matter, gray matter, and CSF partitions. These segmented images were also normalized to the MNI space. The functional images underwent spatial smoothing using a 6 mm FWHM isotropic Gaussian kernel. To conclude the preprocessing, the data’s time series was filtered with a high-pass setting of 128 s.

### Task-based fMRI analysis

Each run utilized a general linear regression model. For this purpose, the task timing was convolved with the hemodynamic response using AFNI’s ’BLOCK’ convolution for marmosets’ data and SPM12 hemodynamic response function for humans’ data. A unified regressor was created for all task conditions using AFNI’s 3dDeconvolve function for marmosets and SPM12 function for humans. The resulting regression coefficient maps for the marmosets were subsequently registered to the template space utilizing the transformation matrices obtained from the anatomical image registrations. We therefore obtained a regression coefficient map from individual runs for each subject, registered to the NIH marmoset brain atlas^28^ for marmosets, and aligned with the MNI brain standard space for humans. These maps were then subjected to group-level comparison through paired t-tests, using AFNI’s 3dttest++ function, ultimately producing Z-value maps. To identify brain regions exhibiting task-positive and task-negative activations in each study, we contrasted all conditions against the baseline period.

For visualization, the resulting z-value functional maps, specific to each study, were displayed using the Connectome Workbench v1.5.076 on fiducial maps, showcasing both the medial and lateral views of the right hemisphere. Additionally, FSLeyes^53^ was used to present the coronal section.

### Task-positive and task-negative probability maps

To examine activation maps across studies, we computed coincidence maps of task-positive and task-negative activations, thresholded at z=3 for task-positive and z=-1 for task-negative, converted them into probability maps using a custom program in Matlab. The resultant probability maps were visualized for each study via the Connectome Workbench v1.5.0^55^ and FSLeyes applications^53^ on fiducial maps of the medial and lateral view of the right hemisphere. Cortical parcellations of the Paxinos marmoset atlas were overlaid^28^. In addition, probability maps were overlaid on coronal sections of a high-resolution (100 x 100 x 100 um) ex-vivo marmoset brain^55^ which was aligned to the NIH marmoset brain template^28^.

### Resting-state fMRI seed analysis

To evaluate the functional connectivity of the hypothalamus – a subcortical area with robust deactivations during task blocks - we measured its task-independent functional connectivity using resting-state fMRI data from an open-access repository (https://marmosetbrainconnectome.org)^30^. This database contains over 70 hours of resting-state fMRI data from 31 awake marmosets (*Callithrix jacchus*, 8 females; age: 14–115 months; weight: 240–625 g). The data were acquired at the University of Western Ontario (5 animals) on a 9.4T scanner and at the National Institutes of Health (26 animals) using a 7T scanner. We positioned a seed (single voxel) in the hypothalamus region which showed the highest probability of activity during baseline periods (9mm anterior, 0.5 mm right, and 7.5 mm dorsal to the anterior commissure, corresponding to the paraventricular hypothalamus). Following this, functional connectivity maps were downloaded and displayed on fiducial and flat maps using the Connectome Workbench (v1.5.0^55^) aligned to the NIH marmoset brain template^28^. These maps were also overlaid on coronal sections of the marmoset brain connectome T2 template using FSLeyes. Additionally, we displayed functional connectivity maps of areas 32, 10, 25, RPB, hypothalamus, hippocampus, and basomedial amygdala on flatmaps of the right hemisphere.

### Anatomical connectivity of frontal cortices with posterior cingulate cortex

To confirm the structural underpinnings of connectivity between marmoset frontal cortex and posterior cingulate, direct intracortical injections of viral anterograde tracers were compared from the BRAIN/MINDS publicly available marmoset tracer resource (https://dataportal.brainminds.jp/marmoset-tracer-injection)^33^. The anterograde tracer injections into the area bordering area 32 and 32v (Brain/MINDS ID: R01_0072; thy1-tTA 1/TRE-clover 1/TRE-Vamp2mPFC 0.25) x10e12 vg/ml) and into area 8aD (Brain/MINDS ID: R01_0026; thy1-tTA 1/TRE-clover 1/TRE-Vamp2mPFC 1) x10e12 vg/ml) were compared. The full details of these injections are publicly available at: https://dataportal.brainminds.jp/.

## Acknowledgement

Support was provided by the Canadian Institutes of Health Research (FRN 148365, S.E.), the Natural Sciences and Engineering Council of Canada (S.E.), and the Canada First Research Excellence Fund to BrainsCAN. We wish to thank Cheryl Vander Tuin, Whitney Froese, Miranda Bellyou, and Hannah Pettypiece for animal preparation and care and Dr. Alex Li for scanning assistance.

## Notes

### Competing Interest Statement

The authors have declared no competing interest.

